# N-terminal degradation activates the Nlrp1b inflammasome

**DOI:** 10.1101/317826

**Authors:** Ashley J. Chui, Marian C. Okondo, Sahana D. Rao, Kuo Gai, Andrew R. Griswold, Brooke A. Vittimberga, Daniel A. Bachovchin

## Abstract

Intracellular pathogens and danger signals trigger the formation of inflammasomes, which activate inflammatory caspases and induce pyroptotic cell death. The anthrax lethal factor metalloprotease and small molecule DPP8/9 inhibitors both activate the Nlrp1b inflammasome, but the molecular mechanism of Nlrp1b activation is not known. Here, we used genome-wide CRISPR/Cas9 knockout screens to identify genes required for Nlrp1b-mediated pyroptosis, and discovered that lethal factor induces cell death *via* the N-end rule proteasomal degradation pathway. Lethal factor directly cleaves Nlrp1b, which induces the N-end rule-mediated degradation of the Nlrp1b N-terminus and thereby frees the Nlrp1b C-terminus to activate caspase-1. DPP8/9 inhibitors also induce proteasomal degradation of the Nlrp1b N-terminus, but, in contrast, not through the N-end rule pathway. Overall, these data reveal that N-terminal degradation is the common mechanism for activation of this innate immune sensor protein.

**One Sentence Summary:** Proteasome-mediated degradation of the Nlrp1b N-terminus releases the Nlrp1b C-terminus to activate caspase-1 and induce pyroptotic cell death.

## Main Text

Mammals express a diverse array of intracellular pattern-recognition receptors (PRRs) that detect cytoplasmic microbial structures and activities, including bacterial flagellin, double stranded DNA, and pathogen modification of host Rho GTPases (*1, 2*). Upon recognition of their cognate danger signals, several PRRs form large, multiprotein complexes called inflammasomes, which recruit and activate the inflammatory protease caspase-1. Active caspase-1, in turn, cleaves and activates inflammatory cytokines and Gsdmd, which mediates a form of lytic cell death called pyroptosis and stimulates a powerful immune response (*1, 3, 4*).

Mouse Nlrp1b is a member of the nucleotide-binding domain and leucine rich repeat containing (NLR) family of intracellular PRRs that can form an inflammasome (*1, 2, 5*). Anthrax lethal toxin (LT) is the best-characterized activator of the Nlrp1b inflammasome (*5, 6*). LT is a bipartite toxin consisting of lethal factor (LF), a zinc metalloprotease, and protective antigen (PA), a cell binding-protein that transports LF into the host cell cytosol. LF directly cleaves Nlrp1b after lysine 44, and this cleavage is required for inflammasome formation (*7-9*). However, it remains unknown how proteolytic cleavage results in the activation of Nlrp1b. We recently discovered that small molecule inhibitors of the host serine dipeptidases DPP8 and DPP9 (DPP8/9) also activate Nlrp1b (*10-12*). It is not known how DPP8/9 inhibition stimulates formation of the Nlrp1b inflammasome, but, unlike LT-induced activation, it does not involve the direct cleavage of Nlrp1b (*11*). Intriguingly, proteasome inhibitors block both LT-and DPP8/9 inhibitor-induced pyroptosis (*11, 13, 14*), but do not block pyroptosis mediated by other inflammasomes (*14, 15*). Thus, although LT and DPP8/9 inhibitors activate Nlrp1b in different ways, a key component of the Nlrp1b activation mechanism – the degradation of some key protein – appears to be shared between these two stimuli.

Here, we wanted to further investigate the mechanism of Nlrp1b activation. We first performed two genome-wide CRISPR/Cas9 screens in the mouse RAW 264.7 macrophage cell line, which is sensitive to both DPP8/9 inhibitors and anthrax LT (*10, 11, 13*), to identify gene knockouts that provide resistance to the DPP8/9 inhibitor Val-boroPro (VbP) and/or LT (Fig. S1A). The top two most enriched sgRNAs for each gene were used to rank each overall gene’s enrichment by RIGER analysis (Fig. 1A, Fig. S1B,C) (*16, 17*). As expected, *Nlrp1b, Casp1,* and *Gsdmd* were among the most enriched genes after both LT and VbP selection. In fact, these were the three highest scoring genes in the VbP screen. *Nlrp1a*, which is not expressed in Balb/c macrophages from which RAW 264.7 cells were derived (*18*), was also enriched, but this is likely due to off-target knockout of *Nlrp1b* by the *Nlrp1a* sgRNAs. Also as expected, many genes that encode proteins known to be required for LT cell penetrance, including the anthrax toxin receptor *Antxr2* (*19*), the cell surface protease furin (*20, 21*), and members of the vacuolar ATPase proton pump (*22*), were highly enriched in LT-treated samples (Fig. 1A).

**Fig. 1.**
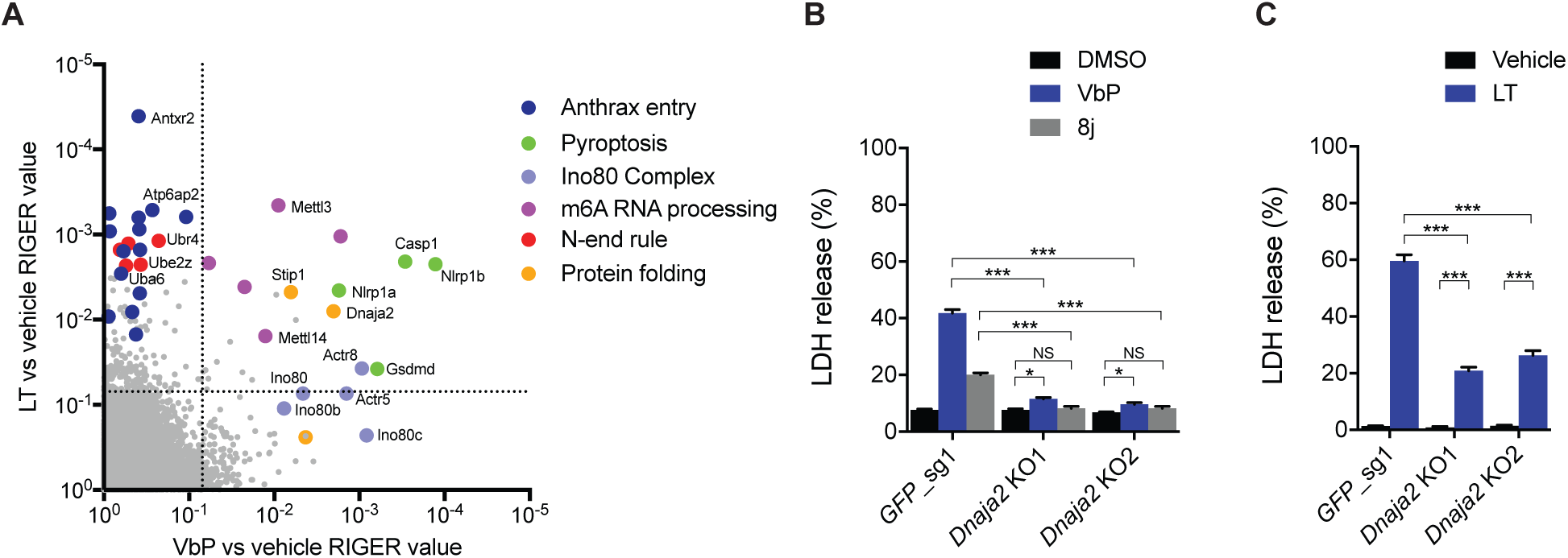
Genome-wide CRISPR/Cas9 screening identifies genes involved in Nlrp1b-mediated pyroptosis. (**A**) RIGER values indicating the relative enrichment of genes after treatment with VbP (x-axis) or LT (y-axis) relative to control. RIGER analysis was performed using the two most enriched sgRNAs for each gene. The dotted lines indicate a RIGER p-value of 0.01. (**B**, **C**) Control or *Dnaja2* KO RAW 264.7 macrophages were treated with VbP (2 μM, 24 h) or 8j (20 μM, 24 h) (**B**) or LT (1 μg/mL, 6 h) (**C**) before cytotoxicity was determined by LDH release. Data are means ± SEM of three biological replicates. ***p < 0.001; *p < 0.05; NS, not significant by two-sided Student’s *t*-test.

In addition to these expected results, several genes with no known association with inflammasome biology were also identified. For example, genes encoding members of the protein folding and chaperoning machinery (*Dnaja2, Stip1,* and *Hsp90ab1)* (*23-25*), the Mettl3-Mettl14 complex that mediates N^6^-adenosine methylation (*Mettl3, Mettl14, Wtap, Zc3h13,* and *Virma*) (*26-28*), and the Ino80 chromatin remodeling complex (*Ino80*, *Ino80b*, *Ino80c*, *Actr5*, and *Actr8*) (*29, 30*) were enriched by LT and/or VbP selection (Fig. 1A). We next wanted to verify a few of these unexpected hits. Indeed, we found that RAW 264.7 cells lacking Dnaja2 (Fig. S2A), a J protein that acts as a co-chaperone of Hsp70s (*23, 31*), were resistant to both DPP8/9 inhibitor-and LT-induced killing (Fig. 1B,C). Treatment of RAW 264.7 cells with sgRNAs targeting *Actr5*, a gene that encodes a core component of the Ino80 complex, did not result in significant Actr5 protein loss (Fig. S2B). However, when cells expressing sgRNAs targeting *Actr5* were treated with VbP, nearly complete Actr5 protein loss was observed (Fig. S2C). These results are consistent with Actr5 deficiency engendering resistance to VbP (Fig. S2D). *Actr5*^-/-^ RAW 264.7 cells proliferated more slowly than control cells (Fig. S2E), accounting for the lack of protein depletion prior to VbP selection. The Ino80 complex genes, including Actr5, were not as enriched in the genome-wide screen by LT as they were by VbP (Fig. 1A), and *Actr5*^-/-^ RAW 264.7 cells were not significantly resistant to LT (Fig. S2F). Interestingly, we observed loss of other Ino80 subunits in *Actr5*^-/-^ cells (Fig. S2C), indicating that loss of one Ino80 subunit affects the stability or expression of the other subunits. Overall, these results confirm that the VbP and LT resistance screens successfully identified many genes, both expected and unexpected, involved in Nlrp1b-dependent pyroptosis.

We were particularly interested in whether any of the newly identified genes could further illuminate the mechanism of Nlrp1b inflammasome activation, and, in particular, the role of the proteasome. On this note, we noticed that several genes involved in the N-end rule proteasomal degradation pathway (*32-34*), *Ubr2, Ubr4, Uba6, Ube2z,* and *Kcmf1,* were highly enriched by LT but not by VbP (Fig. 1A, Fig. S1C). The N-end rule pathway recognizes, ubiquitinates, and degrades proteins with destabilizing N-terminal residues (*32, 33*). Interestingly, Wickliffe and co-workers showed that inhibitors of the N-end rule pathway, bestatin methyl ester and amino acid derivatives (e.g., L-Phe-NH_2_ and L-Trp-NH_2_), block LT-mediated cell death (*35*). Based on these data, the authors proposed that the LF protease might cleave a key substrate protein to generate a destabilizing N-terminal residue, inducing that protein’s degradation *via* the N-end rule and triggering cell death. However, the key protein substrate of possible N-end rule degradation was not identified, and the direct involvement of N-end rule proteins was not established. Our screening results strongly indicate that the N-end rule pathway is indeed involved in LT-mediated cytotoxicity. We hypothesized that Nlrp1b itself, which was discovered to be directly cleaved by LF after the Wickliffe study (*7-9*), might be the key LF substrate that is degraded by the N-end rule pathway.

The Nlrp1b protein, like other NLR proteins, contains nucleotide-binding (NACHT) and leucine-rich repeat (LRR) domains (Fig. 2A). However, unlike other NLR proteins, Nlrp1b has C-terminal “function-to-find” (FIIND) and caspase activation and recruitment (CARD) domains. Nlrp1b undergoes post-translational autoproteolysis within the FIIND domain, resulting in N-and C-terminal fragments that remain associated in an auto-inhibited state (*36-38*). Autoproteolysis is necessary for inflammasome formation (*11, 37, 38*), but why autoproteolysis is necessary remains unknown. We first wanted to confirm which domains of Nlrp1b were required for cytotoxicity. We transfected HEK 293T cells stably expressing mouse caspase-1 with constructs encoding various fragments of Nlrp1b (Fig. 2B,C). We found that the N-terminal fragment preceding the FIIND autoproteolysis site was completely non-toxic, but all of the C-terminal constructs containing the CARD were toxic. Although the CARD alone did induce some toxicity, maximal toxicity required at least a small portion of the FIIND domain (~50 amino acids), consistent with a previous report (*39*). Based on these results, we predicted that N-end rule degradation of the Nlrp1b N-terminus could potentially free the C-terminus to activate caspase-1, as the break in the polypeptide chain in the FIIND domain would prevent the concomitant destruction of the C-terminal CARD domain.

**Fig. 2.**
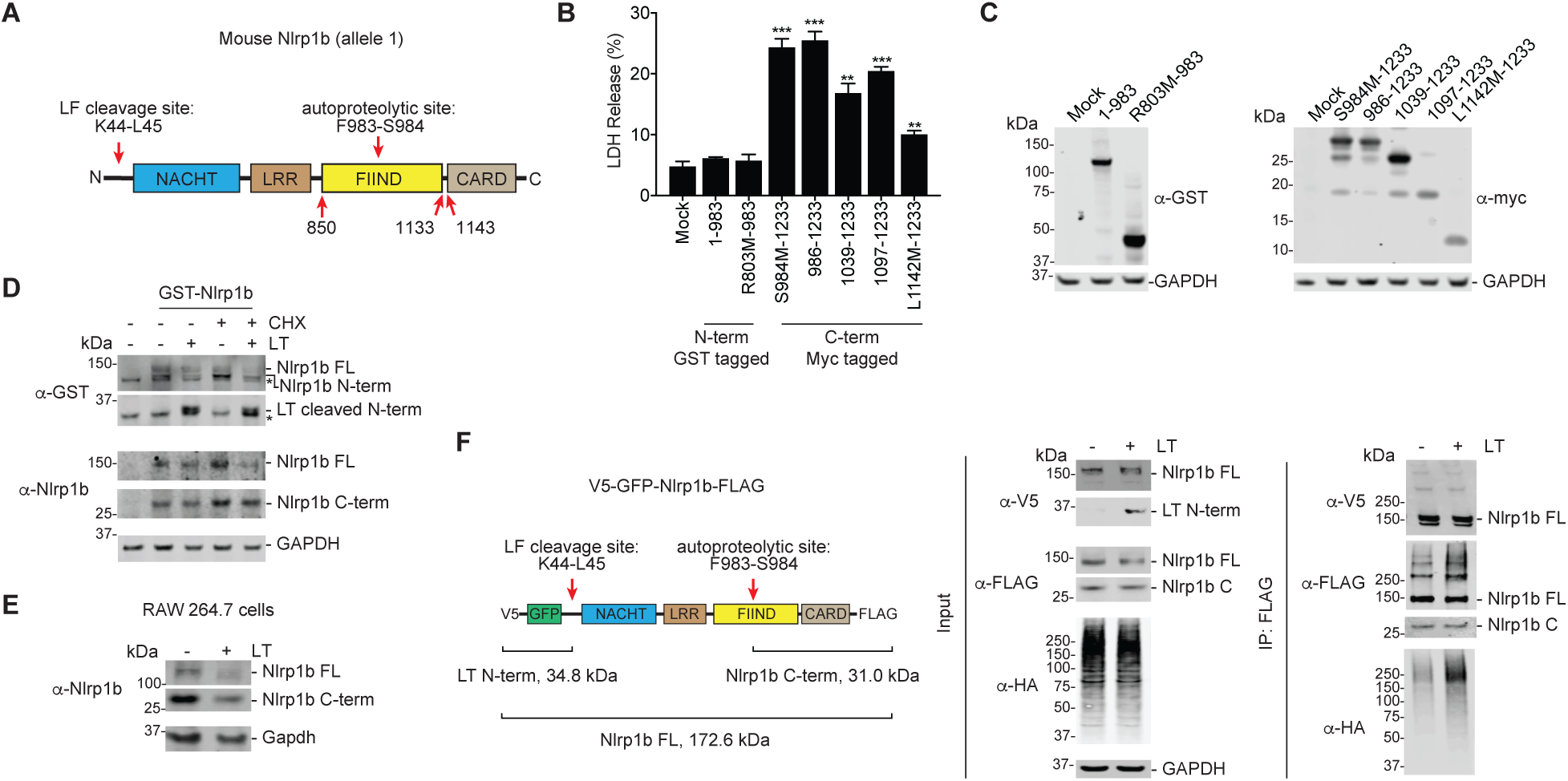
LT induces Nlrp1b ubiquitination and degradation. (**A**) Diagram of mouse Nlrp1b (allele 1). The LF cleavage and FIIND autoproteolysis sites are indicated. The cartoon is not drawn to scale. (**B,C**) HEK 293T cells stably expressing mCasp1 were transiently transfected with the indicated constructs (2 μg) for 24 hours, before cell viability was evaluated by LDH release (**B**) and expression was evaluated by immunoblotting (**C**). The N-terminal constructs had N-terminal GST tags and the C-terminal constructs had C-terminal Myc-tags. Residues that were mutated to create start sites are indicated. Data are means ± SEM of three biological replicates. ***p < 0.001, **p < 0.01 by two-sided Student’s *t*-test. (**D**) HEK 293T cells stably expressing mCasp1 were transiently transfected with a construct encoding GST-Nlrp1b (0.03 μg). After 24h, cycloheximide (100 μg/mL) and/or LT (1 μg/mL) was added to the indicated samples, which were then incubated for an additional 6 h. Protein levels were evaluated by immunoblotting. (**E**) RAW 264.7 cells were treated with LT (1 μg/mL, 6 h) before protein levels were evaluated by immunoblotting. (**F**) Constructs encoding V5-GFP-Nlrp1b-FLAG (shown) and HA-ubiquitin were co-transfected into HEK 293T cells. After 24 hours, cells were treated with bortezomib (1 μg/mL) and BSA (1 μg/mL) or LT (1 μg/mL) for 3 h before lysates were harvested, immunoprecipitated with anti-FLAG beads, and evaluated by immunoblotting.

We next asked whether the Nlrp1b protein was degraded after LT cleavage. HEK 293T cells ectopically expressing caspase-1 and Nlrp1b are sensitive to LT-and VbP-induced pyroptosis (*11*). Consistent with our hypothesis, we saw loss of the full-length Nlrp1b protein after treatment of these cells with LT (Fig. 2D). Nlrp1b protein loss was even more pronounced when the cells were pre-treated with cycloheximide to block new protein synthesis. It should be noted that a loss of GST detection in this assay could be simply due to LF proteolytic cleavage and removal of the N-terminal GST-tag. However, the loss of direct Nlrp1b detection (the antibody detects the C-terminus) without the appearance of a slightly lower band is strongly indicative of degradation. Consistent with our proposed model, we saw less loss of the C-terminal fragment relative to the full-length protein, although some of the C-terminal fragment was also degraded. Similar degradation of the endogenous Nlrp1b protein was observed in RAW 264.7 cells after LT treatment (Fig. 2E). Interestingly, VbP also triggered the degradation of Nlrp1b (Fig. S3), indicating that the degradation mechanism of activation is shared between the two stimuli.

We next wanted to determine if LT induces the ubiquitination of Nlrp1b. We transfected Nlrp1b with an N-terminal V5-GFP tag and a C-terminal FLAG tag together with HA-tagged ubiquitin in HEK 293T cells (Fig. 2F). After 24 hours, cells were treated with LT and bortezomib (to block degradation of ubiquitinated proteins) for 3 h before lysates were harvested. Some, but not all, Nlrp1b was cleaved by LT at this time point (Fig. 2F). Nlrp1b was then immunoprecipitated using anti-FLAG beads and probed with V5, FLAG, and HA antibodies. Consistent with LT-induced ubiquitination of Nlrp1b, we observed greater amounts of HA-tagged and high molecular weight FLAG-tagged species in LT-treated lysates. Notably, we did not see a greater amount of high molecular weight V5-tagged species, which is consistent with the LF-uncleaved protein not being ubiquitinated.

We next wanted to confirm that the N-end rule pathway mediated LT-induced degradation of Nlrp1b and was required for LT-induced pyroptosis. Bestatin methyl ester is a cell permeable analog of bestatin, a non-specific inhibitor of aminopeptidases. Aminopeptidases remove N-terminal amino acids from polypeptides, and therefore can reveal destabilizing residues and induce protein degradation *via* the N-end rule pathway. As such, bestatin methyl ester can inhibit N-end rule-mediated protein degradation (*35*). As expected, bestatin methyl ester and bortezomib block LT-induced pyroptosis in RAW 264.7 cells (Fig. 3A) (*35*), and both completely inhibit the degradation of Nlrp1b induced by LT in these cells (Fig. 3B). To confirm that N-end rule proteins were involved in LT-induced cell death, we treated RAW 264.7 cells with sgRNAs targeting *Ubr2*, which encodes an N-end rule E3 ligase (*32, 33*) that was highly enriched in the LT screen (Fig. 3C). LT induced considerable Nlrp1b degradation in control, but not in Ubr2 deficient, cells at 3 h (Fig. 3C). Some LT-induced degradation of Nlrp1b was observed in Ubr2 deficient cells at 6 h (Fig. 3C), but the amount of LT-induced cell death was still significantly reduced (Fig. 3D). That Ubr2 deficient cells are not completely resistant to LT is likely due to the redundancy between the N-end rule E3 ligases (for example, the N-end rule E3 ligase Ubr4 was also highly enriched in the LT screen). Regardless, these data confirm that the N-end rule pathway mediates LT-induced Nlrp1b degradation and cell death. As expected, loss of Ubr2 did not block VbP-induced pyroptosis (Fig. 3E,F, Fig. S4)

**Fig. 3.**
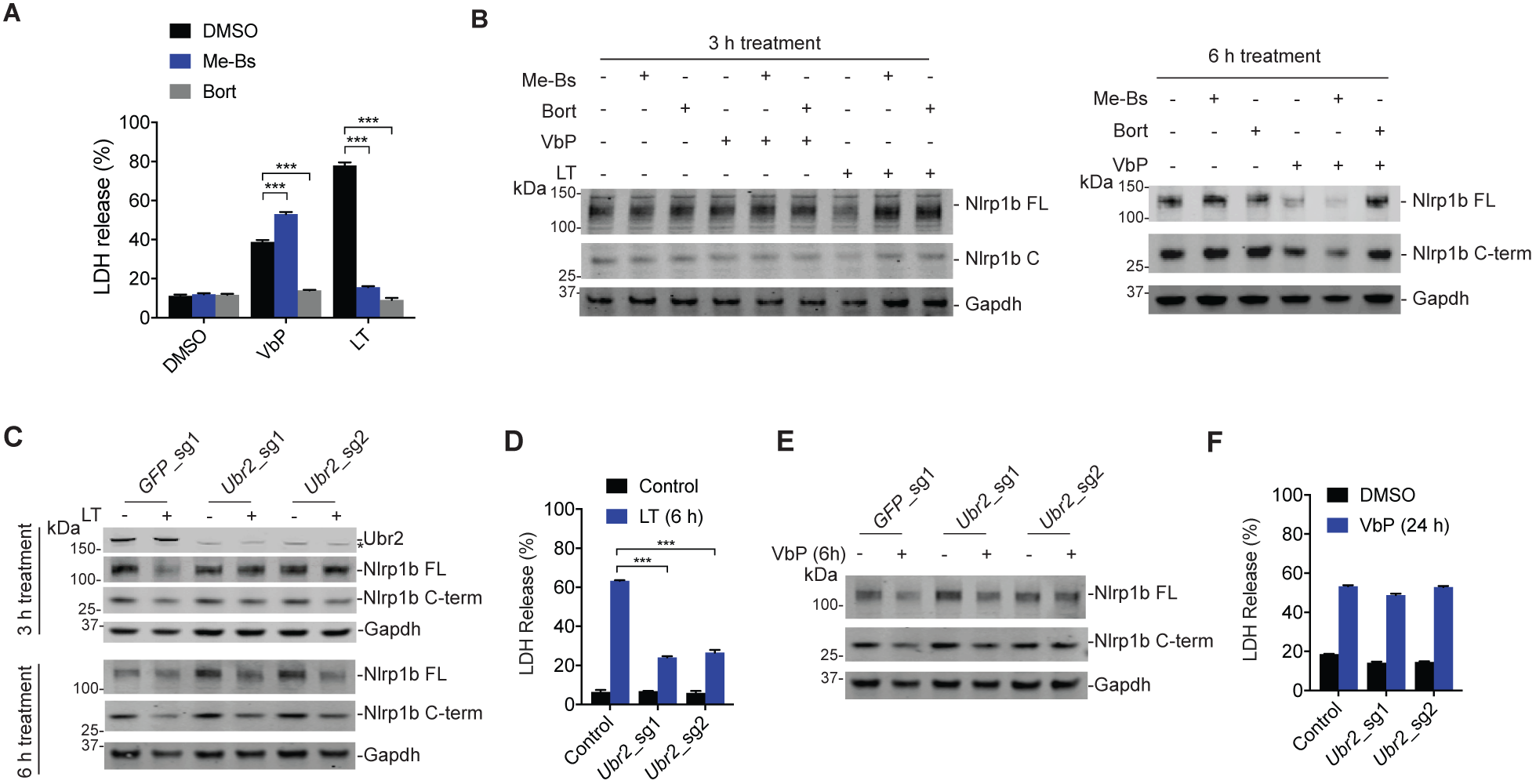
The N-end rule pathway mediates LT-induced Nlrp1b degradation. (**A**) RAW 264.7 cells were treated with the DMSO, VbP (2 μM), or LT (1 μg/mL) and either bestatin methyl ester (Me-Bs, 10 μM) or bortezomib (bort, μM) for 6 h before supernatants were evaluated for LDH release. Asterisk indicates a background band. (**B**) RAW 264.7 cells were treated with the indicated agents (concentrations as in **A**) for 3 h (left panel) or 6 h (right panel) before lysates were harvested and evaluated for protein levels by immunoblotting. (**C-F**) RAW 264.7 cells treated with the indicated sgRNAs were incubated with VbP (10 μM) or LT (1 μg/mL) for the indicated time intervals. In **C** and **E**, lysates were probed for protein levels by immunoblotting. In **D** and **F**, cytotoxicity was evaluated by LDH release. A 6 h time point for LDH release by VbP is shown in Fig. S4. In **A**, **D**, and **F** data are means ± SEM of three biological replicates. ***p < 0.001 by two-sided Student’s *t*-test.

Similarly, bortezomib blocks VbP-induced cell death and VbP-induced degradation of Nlrp1b in RAW 264.7 cells (Fig. 3A,B). It should be noted that VbP-induced cell death is markedly slower than LT-induced cell death in RAW 264.7 cells (LT induces significant cell death at ~ 3 h, whereas VbP requires ~6 h) (*11*), and correspondingly VbP-induced Nlrp1b degradation occurs more slowly (Fig. 3B). We predicted that bestatin methyl ester would have no effect on VbP-induced cell death or Nlrp1b degradation. However, we unexpectedly found that bestatin methyl ester actually potentiated VbP-induced cell death (Fig. 3A) and Nlrp1b degradation (Fig. 3B). This effect was not an artifact of the RAW 264.7 cell line, as bestatin and bestatin methyl ester increased the levels of G-CSF induced by VbP in C57BL/6J mice (Fig. 4A) and augmented DPP8/9 inhibitor-induced cell killing in human THP-1 cells (Fig. S5). We reasoned that aminopeptidase inhibition might be increasing the levels of the pyroptosis-inducing DPP8/9 substrate (i.e., aminopeptidases could cleave and inactivate this putative substrate in the absence of DPP8/9 activity), or that the N-end rule pathway also degrades and inactivates the C-terminus of Nlrp1b (i.e., inhibition of the N-end rule could enhance FIIND-CARD toxicity). The former possibility seemed most likely, as bestatin methyl ester enhanced VbP-induced Nlrp1b degradation (Fig. 3B). To rule out any involvement of the latter possibility, we expressed the FIIND-CARD fragment of Nlrp1b with an S984M start site (Fig. 4B), or fused in frame to ubiquitin to generate the native S984 start site (Fig. 4C), and treated with agents that either activate or inhibit this pyroptotic pathway. These agents had no effect on the toxicity of the FIIND-CARD fragment, indicating that all regulation occurs at the level of or upstream of the full-length Nlrp1b protein containing the N-terminus.

**Fig. 4.**
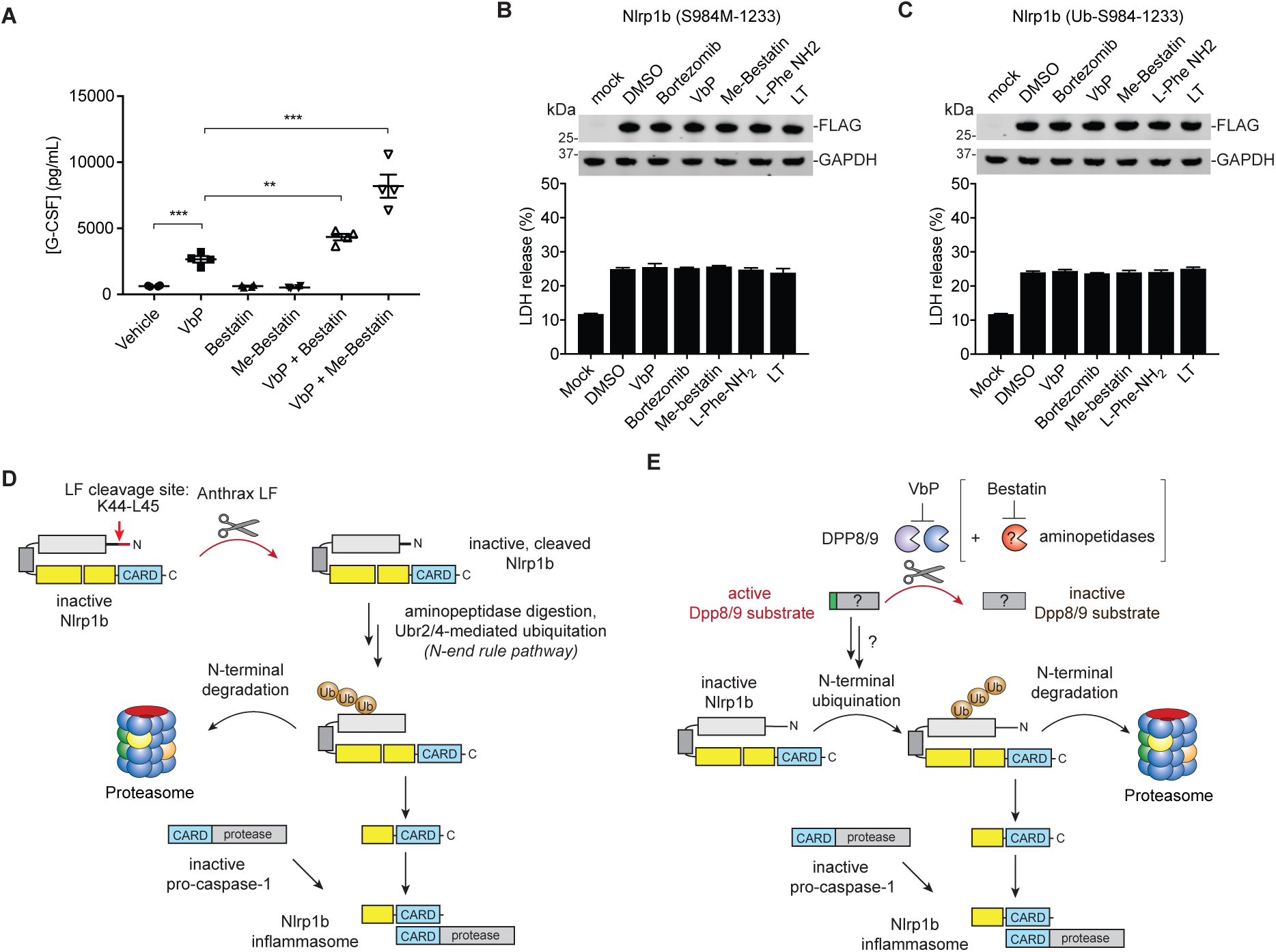
The toxicity of the Nlrp1b C-terminus is not affected by N-end rule, DPP8/9, or proteasome inhibition. (**A**) C57BL/6J mice were dosed intraperitoneally with vehicle (0.1 N HCl), VbP (2 μg/mouse), bestatin (50 mg/kg), bestatin methyl ester (50 mg/kg), or the indicated combinations. After 6 h, levels of serum G-CSF were determined by ELISA. Data are means ± SEM, n = 4 mice/group. **p < 0.01, ***p < 0.001 by two-sided Student’s *t*-test. (**B,C**) HEK 293T cells stably expressing mouse caspase-1 were transiently transfected with constructs (0.2 μg) encoding the FIIND-CARD fragment of Nlrp1b with an S984M start site (**B**) or fused in frame to ubiquitin to generate the native S984 start site (**C**). After 24 h, cells were treated with the indicated agents for 6 h before supernatants were evaluated for LDH release. Data are means ± SEM of three biological replicates. (**D**,**E**) Proposed models of Nlrp1b activation by LT (**D**) and VbP (**E**). Bestatin is in brackets because the precise role of its aminopeptidase targets upstream of Nlrp1b has not been established.

Overall, this work has revealed that N-terminal degradation is a common mechanism of Nlrp1b inflammasome activation. Intriguingly, there are at least two distinct mechanisms, one direct (Fig. 4D) and one indirect (Fig. 4E), that mediate N-terminal degradation. In the direct mechanism identified here, LF protease cleaves Nlrp1b and generates a destabilizing N-terminal residue. As it is unlikely that anthrax LF evolved to intentionally trigger pyroptosis, we speculate that Nlrp1b may serve as a molecular ‘booby trap’ for LF and possibly other pathogen encoded activities. Consistent with this premise, the Vance laboratory has recently discovered that the *S. flexneri* E3 ligase IpaH7.8 ubiquitinates and degrades Nlrp1b to activate this inflammasome (Sandstrom *et al* (2018) *biorxiv.org*). In the indirect mechanism identified here, we found that the inhibition of DPP8/9 activates an endogenous proteasomal degradation pathway to destroy the Nlrp1b N-terminus. The molecular details of this pathway remain to be determined, and it is not yet clear if this mechanism evolved to sense a specific pathogen or danger signal or if it serves some other purpose. Regardless, here we have identified many genes critical for Nlrp1b-mediated pyroptosis and have demonstrated that N-terminal degradation is a unifying mechanism for Nlrp1b activation. We expect that future studies will leverage these insights to further clarify how Nlrp1b senses specific pathogens and how this inflammasome can be modulated for therapeutic benefit.

## Acknowledgments

We thank W. Bachovchin, W. Wu, and J. Lai for Val-boroPro and 8j, the Vance laboratory for the Nlrp1b antibody, and R. Garippa for assistance with the genome-wide screen.

## Author contributions

D.A.B. conceived and directed the project, performed experiments, and analyzed data, and wrote the paper; A.J.C, M.C.O., S.D.R., K.G., A.R.G, and B.A.V. performed experiments and analyzed data.

## Data and materials availability

All data is available in the main text or the supplementary materials.

## Materials and Methods

### Cloning

sgRNAs were designed using the Broad Institute’s web portal (*40*) (http://www.broadinstitute.org/rnai/public/analysis-tools/sgrna-design) and cloned into the lentiGuide-puro vector (Addgene #52963) as described previously (*41*). The sgRNA sequences are listed in **Table S1**. cDNA encoding the full-length mouse *Nlrp1b* gene was cloned from RAW 264.7 macrophages and moved into the pDEST27 vector (Addgene) or a modified pLEX_307 vector (Addgene) with an N-terminal V5-GFP tag and C-terminal FLAG tag using Gateway technology (Thermo Fisher Scientific). The indicated m*Nlrp1b* fragments were subcloned into the pDEST27 vector or into modified pLEX_307 vectors with a C-terminal FLAG or Myc tags. cDNA encoding mouse *Casp1* was purchased from Origene and was subcloned into a modified pLEX_307 vector with a hygromycin resistance marker. A vector encoding HA-ubiquitin was purchased from Addgene (#18712).

### Reagents and antibodies

Val-boroPro (*42*) and compound 8j (*43*) were synthesized according to previously published protocols. For cell culture experiments, Val-boroPro was resuspended in DMSO containing 0.1% TFA to prevent compound cyclization. Anthrax lethal toxin was purchased from List Biological Laboratories, bortezomib from LC laboratories, bestatin methyl ester, L-phenylalaninamide hydrochloride, amastatin, and cycloheximide from Sigma-Aldrich, CHR-2797 from Tocris Bioscience, batamistat from R&D Systems, bestatin from VWR, and actinonin from Enzo Life Sciences. Antibodies used include: mouse caspase-1 (clone Casper-1, Adipogen), GAPDH (clone 14C10, Cell Signaling Technology), V5 (ab9116, Abcam), GST (26H1, Cell Signaling Technology), Myc (9B11, Cell Signaling Technology), FLAG M2 (F1804, Sigma-Aldrich), HA (ab9110, Abcam), Ubr2 (ab217069, Abcam), Dnaja2 (ab157216, Abcam), Actr5 (21505-1-AP, Proteintech), Ino80b (ab175117, Abcam), Actr8 (ab177335, Abcam), PARP (#9542, Cell Signaling Technology), GSDMD (NBP2-33422, Novus Biologicals), and Nlrp1b (this monoclonal antibody was a gift from the Vance laboratory at UC Berkeley).

### Cell culture

HEK 293T and RAW 264.7 cells were purchased from ATCC and grown in Dulbecco’s modified Eagle’s medium (DMEM) with 10% fetal bovine serum (FBS). THP-1 cells were purchased from ATCC and grown in RPMI-1640 medium with 10% FBS. All cells were grown at 37 °C in a 5% CO_2_ incubator. Cell lines were regularly tested for mycoplasma using the MycoAlert^TM^ Mycoplasma Detection Kit (Lonza).

### LDH cytotoxicity assays

RAW 264.7 cells were seeded at 0.5 × 10^6^ cells/well in 6-well plates in standard growth medium 24 hours prior to treatment. Cells were treated with the indicated agents for the indicated times. Supernatants were then harvested and analyzed for LDH activity using an LDH cytotoxicity assay kit (Pierce). For experiments with proteasome inhibitors or bestatin methyl ester, cells were treated with these agents 30 minutes prior to the addition of Val-boroPro and/or LT, and then incubated for the indicated amount of time. For LDH release experiments in HEK 293T cells, we first generated cells stably expressing mCasp1. HEK 293T cells were transfected with 2 μg of the pLEX_307 containing *mCasp1*, 2 μg PAX2 (Addgene, #35002), and 1 μg pMD2.G (Addgene, #12259) using the Fugene HD transfection reagent (Promega) and incubated for 48 hours. HEK 293T cells were then spinfected with virus-containing supernatant from the transfection for 2 hours at 1000 x *g* at 30 °C supplemented with 8 μg/mL polybrene. After 2 days, cells were selected for stable expression of mCasp1 using hygromycin (100 μg/mL). HEK 293T cells expressing mCasp1 were then seeded at 0.5 × 10^6^ cells/well in 6-well plates in standard growth medium, and 24 h later transiently transfected with indicated constructs encoding *Nlrp1b* fragments using Fugene HD. After 24 hours, supernatant was then harvested and analyzed for LDH activity, or cells were treated with the indicated agents for 6 additional h before LDH release was assessed.

### CellTiter-Glo cell viability assay

*DPP8/9*^-/-^ and *CASP1*^-/-^ THP-1 cells were generated previously (*10*). THP-1 and RAW 264.7 cells were plated (2000 cells/well) in white, 384-well clear bottom plates (Corning) using an EL406 Microplate Washer/Dispenser (BioTek) in 25 μL final volume of media. Compounds were added to THP-1 cells using a pintool (CyBio). THP-1 cells were incubated for 48 hours at 37 °C, and RAW 264.7 cells were incubated for the indicated times at 37 °C. Assay plates were then removed from the incubator and allowed to equilibrate to room temperature on the bench top before addition of 10 μL of CellTiter-Glo reagent (Promega) according to the manufacturer’s instructions. Assay plates were shaken on an orbital shaker for 2 minutes and incubated at room temperature on the bench top for 10 minutes. Luminescence was then read using a Cytation 5 Cell Imaging Multi-Mode Reader (BioTek).

### Knockout cell lines

RAW 264.7 cells stably expressing Cas9 were generated previously (*10*). Constructs encoding sgRNAs were packaged into lentivirus in HEK 293T cells using the Fugene HD transfection reagent (Promega) and 2 μg of the vector, μ2 g PAX2 (Addgene, #35002), and 1 μg pMD2.G (Addgene, #12259), and incubated for 48 h. RAW 264.7 cells were spinfected with virus-containing supernatant from the transfection for 2 hours at 1000 × *g* at 30 °C without polybrene. After 2 days, cells were selected for stable expression of sgRNAs using puromycin (5 μg/mL). After 10 days, single cells were isolated by serial dilution and expanded to obtain complete knockouts. RAW 264.7 cells treated with sgRNAs targeting *Actr5* were incubated with VbP (10 μM) for 1 week to obtain Actr5-deficient cells.

### Cytokine stimulation in mice

For cytokine induction in C57BL/6J mice (The Jackson Laboratory), 9-week-old male mice were treated intraperitoneally with 100 μL of vehicle (1 N HCl in PBS, pH = 7.4), 100 μL of Val-boroPro (20 μg/ 100 μL), 100 μL of bestatin (50 mg/kg), 100 μL of bestatin methyl ester (50 mg/kg), or the indicated combinations. Val-boroPro was stored at 10x final concentration in 0.01N HCl and diluted into PBS immediately before dosing. Bestatin and bestatin methyl ester was stored at 10x final concentration in DMSO and diluted into PBS immediately before dosing. Serum was collected 6 h after dosing via retro-orbital collection and G-CSF levels were measured by Quantikine ELISA (R&D Systems). This animal protocol was reviewed and approved by the Memorial Sloan Kettering Cancer Center Institutional Animal Care and Use Committee (IACUC). Sample size was based on the statistical analysis of previous experiments with vehicle treated mice versus Val-boroPro treated mice (*10, 44*). No animals were excluded. The experiments were not randomized and the investigators were not blinded.

### Immunoblotting experiments

RAW 264.7 macrophages were seeded at 0.5 × 10^6^ cells/well in 6-well plates or at 5 × 10^6^ cells/dish in 10 cm plates, and 24 h later were treated with agents as described. Bestatin methyl ester and bortezomib, if used, were added 30 minutes prior to LT or VbP treatment. For experiments using HEK 293T cells stably expressing mCasp1, cells were seeded at 0.5 × 10^6^ cells/well in 6-well plates, and 24 h later transiently transfected with the indicated constructs. After 24 h, lysates were then harvested, or cells treated with agents as indicated before lysates were harvested. Lysates were normalized to 1 mg/mL using the DC Protein Assay kit (Bio-Rad), separated by SDS-PAGE, immunoblotted, and visualized using the Odyssey Imaging System (Li-Cor).

### Immunoprecipitation experiments

For immunoprecipitation experiments, HEK 293T cells were seeded at 7 × 10^6^ cells/well in 10 cm plates and transiently transfected with 5 μg of N-terminal V5 and C-terminal FLAG-tagged Nlrp1b and 5 μg of N-terminal HA-tagged ubiquitin using Fugene HD. After 24 h, cells were then treated with bortezomib for 30 minutes followed by LT or vehicle for an additional 3 hours and harvested. Cells were lysed by sonication and normalized to 1 mg/mL using the DC Protein Assay kit (Bio-Rad). Immunoprecipitations were carried out by incubating 1 mg total cell lysate prepared in PBS with protease inhibitors (Halt Protease Inhibitor Cocktail, Thermo Fisher) and 40 μL of anti-FLAG-M2 agarose resin (Sigma-Aldrich) in 1 mL total volume overnight at 4 °C. Beads were then washed 3X in PBS (pH 7.4), and bound proteins were eluted in 3X FLAG peptide (150 ng/μL). The samples were separated by SDS-PAGE, immunoblotted, and visualized using the Odyssey Imaging System (Li-Cor).

### Genome wide CRISPR-Cas9 Knockout Screens

*Lentiviral Production:* The mouse GeCKO 2-vector system sgRNA pool B library (Addgene, # 1000000053) was amplified and verified by the RNAi core at MSKCC following established protocols (*45*). For lentiviral production, HEK 293T cells were seeded at 15 x10^6^ cells in 4 x 15 cm dishes in DMEM with 10% FBS. 24 h later, cells were transfected with 15 μg Pax2 and 10 μg pMD2.G (Addgene # 35002 and 12259, respectively) using 75 μl of Fugene HD reagent in 1.5 mL OPTI-MEM. Media was replaced 6-8 h after transfection with DMEM containing 30% FBS. 2 days later, lentivirus was harvested, aliquoted and frozen at -80 °C. Viral titer was measured following established protocols (*16*). Briefly, RAW 264.7 cells stably expressing Cas9 (*10*) were seeded at 3 x 10^6^ cells/well in 6-well plates. A range of virus-containing supernatant (0-1000uL) was added and the multiplicity of infection (MOI) was determined by calculating the number of transduced cells 3 days after selection with puromycin. *VbP and LT selection screens*: 180 x 10^6^ RAW 264.7 cells stably expressing Cas 9 were seeded at 3 x 10^6^ cells/well in 6 well plates. The pooled library was transduced at an MOI of 0.3 with a coverage of >500 cells per gRNA. Plates were spinfected at 1500*g* for 90 min and incubated for 72 h. Cells were then selected with puromycin (5 μg/ml) for 14 days. The transduced cells were seeded at 50 x 10^6^ cells for each replicate of control, VbP (2 μM) and LT (1μg/mL) treatments (n=3/treatment). The untreated cells were harvested 24h after seeding. VbP treated cells were treated with drug for 1 week before being washed 3 x with PBS (10 mL) and propagated for an additional 5 days before harvesting. For LT selection, cells were treated with 1 μg/ml each of protective antigen (PA) and lethal factor (LF) for 3h. Cells were washed twice with PBS and allowed to propagate for 4 days after replacing media. Cells were then selected similarly with LT for a second time. Cells were harvested at seeding concentration and cell pellets frozen. *Illumina library preparation:* Frozen cell pellets were thawed and genomic DNA was harvested using the DNeasy blood and tissue kit (Qiagen # 69506). A two-step PCR protocol (*16*) with Phusion high fidelity DNA polymerase (New England BioLabs) was performed following the manufactured protocol (NEB M0530). We used established primer sequences for amplifying the Lentiguide puro sgRNA for the first PCR (*16*) and standard Illumina primers and adaptor sequences for the second PCR (Addgene). The amplicons were extracted from 2% agarose gel using the QIAquick Gel Extraction Kit (Qiagen # 28076). The samples were quantified, pooled and sequenced on a Hiseq2500 by the Integrated Genomics Operation at MSKCC. *RIGER analysis:* A pseudocount of one was added to all read counts for each sgRNA in each treatment group. Each sgRNA was normalized to its relative representation in each sample. Fold-enrichment was determined by dividing the average sgRNA representation in the treated samples by the average sgRNA representation in the control samples. RIGER analysis was performed using GENE-E software (https://software.broadinstitute.org/GENE-E/) using the Second Best Rank algorithm that takes into account the combined sum of the first and second best ranks for sgRNAs for a given gene, as described on the GENE-E website.

---

**Fig. S1.**
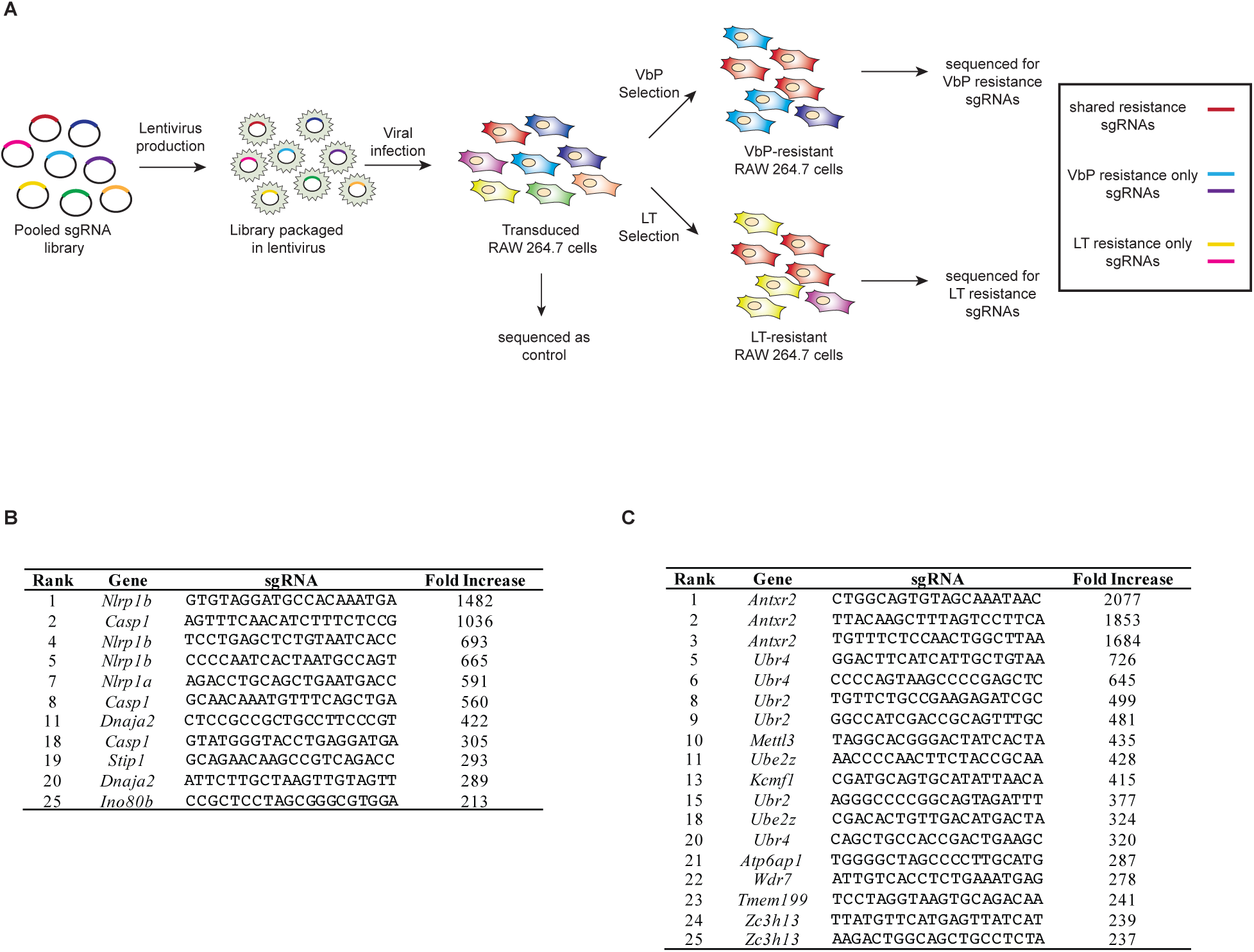
Overview of the CRISPR/Cas9 genome-wide screens for VbP and LT resistance. (**A**) Schematic of the genome-wide screen workflow. (**B**,**C**) sgRNAs for the VbP screen (**B**) and the LT screen (**C**) were ranked by overall fold enrichment relative to the control. Shown are sgRNAs that rank in the top 25 by overall fold enrichment for which additional sgRNAs targeting the same gene also ranked in the top 500 by overall fold enrichment.

**Fig. S2.**
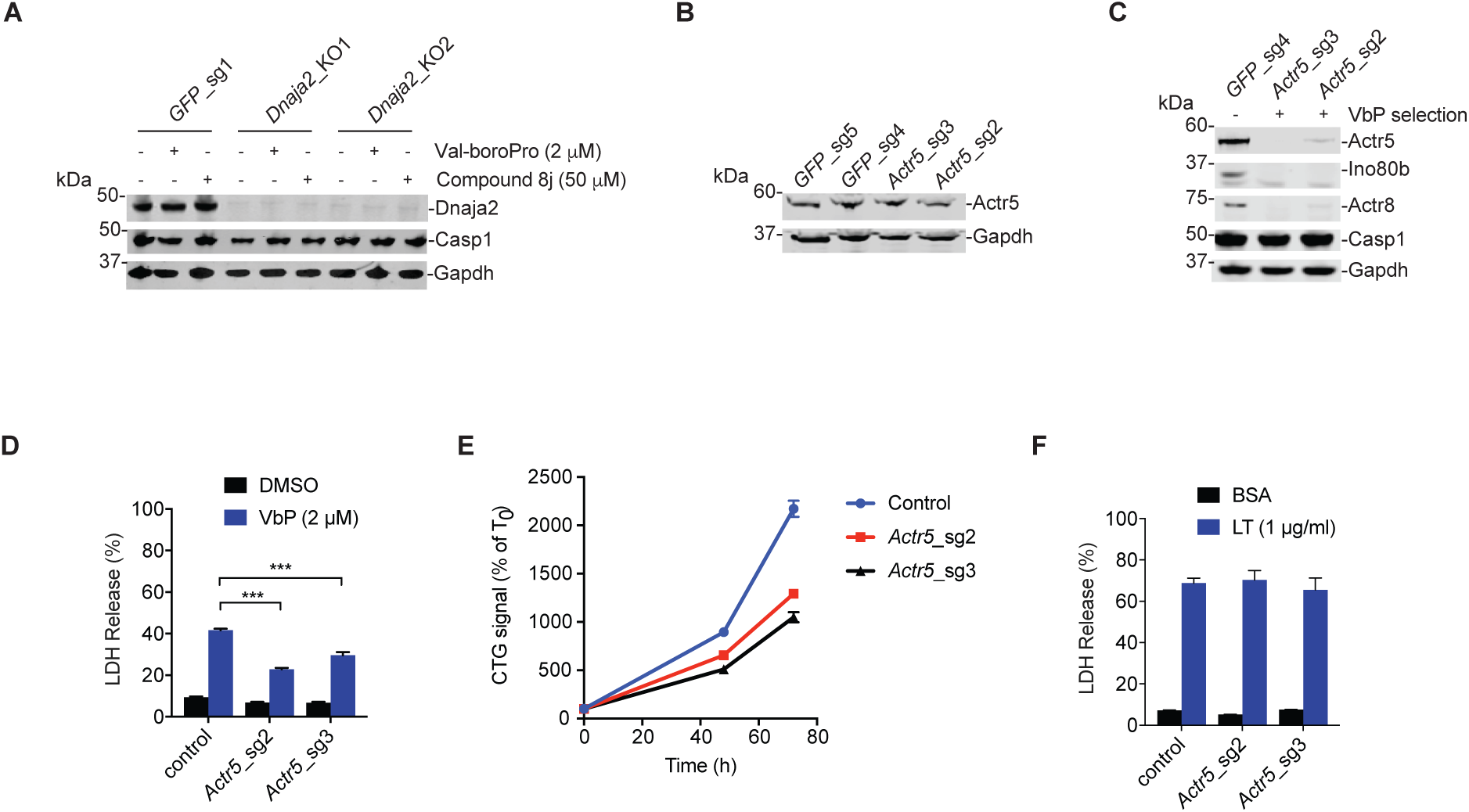
Validation of hits in the genome-wide CRISPR/Cas9 screens. (**A**) Confirmation of *Dnaja2* knockout in RAW 264.7 cells by immunoblotting. Cells were treated with DMSO, VbP, or compound 8j for 24 h prior to immunoblotting. (**B**) Evaluation of Actr5 protein loss after treatment of RAW 264.7 cells with sgRNAs targeting *Actr5* by immunoblotting. (**C**) Evaluation of Actr5, Ino80b, and Actr8 protein levels by immunoblotting after treatment of RAW 264.7 cells in **B** with VbP (10 μM) for 1 week. (**D**) RAW 264.7 macrophages from **C** were treated with VbP (24 h) before cytotoxicity was determined by LDH release. Data are means ± SEM of four biological replicates. ***p < 0.001 by two-sided Student’s *t*-test. (**E**) Cell proliferation time-course of the control and *Actr5* knockout cells from **C** as measured by CellTiter-Glo. Data are means ± SEM of ten biological replicates. (**F**) RAW 264.7 macrophages from **C** were treated with LT (3 h) before cytotoxicity was determined by LDH release. Data are means ± SEM of four biological replicates.

**Fig. S3.**
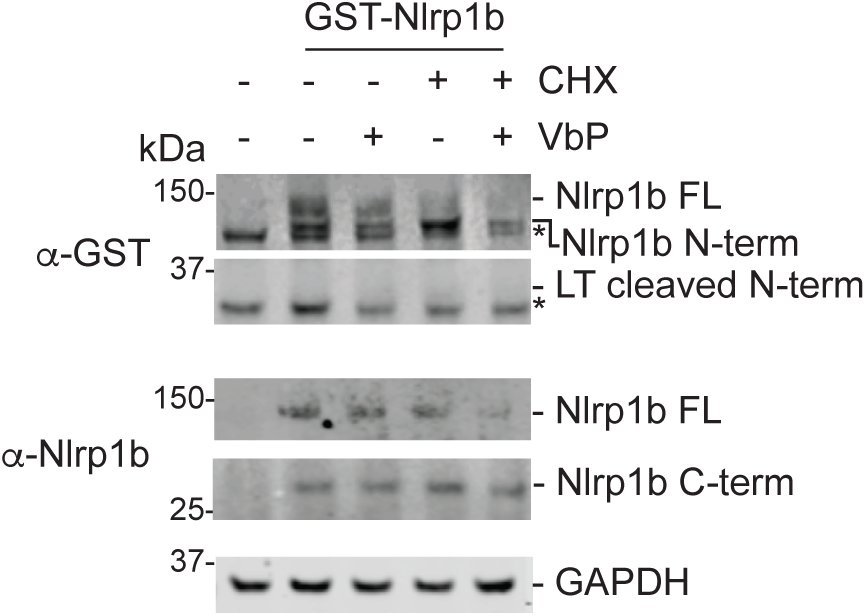
Val-boroPro induces degradation of Nlrp1b. HEK 293T cells stably expressing mouse caspase-1 were transiently transfected with a construct encoding GST-Nlrp1b. After 24h, cycloheximide (100 μg/mL) and/or VbP (2 μM) was added to the indicated samples, which were then incubated for an additional 6 h. Protein levels were then evaluated by immunoblotting. Asterisks indicate background bands. VbP does not induce the formation of the LT cleaved N-terminus, as expected.

**Fig. S4.**
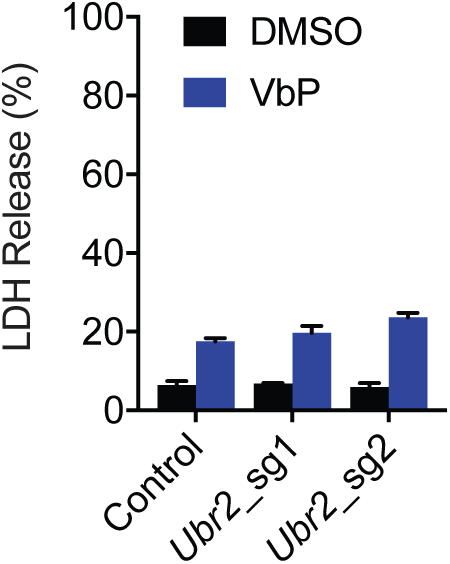
Ubr2 deficiency does not block VbP-induced pyroptosis. RAW 264.7 cells treated with the indicated sgRNAs were incubated with VbP (10 μM) for 6h before cytotoxicity was evaluated by LDH release. Data are means ± SEM of three biological replicates.

**Fig. S5.**
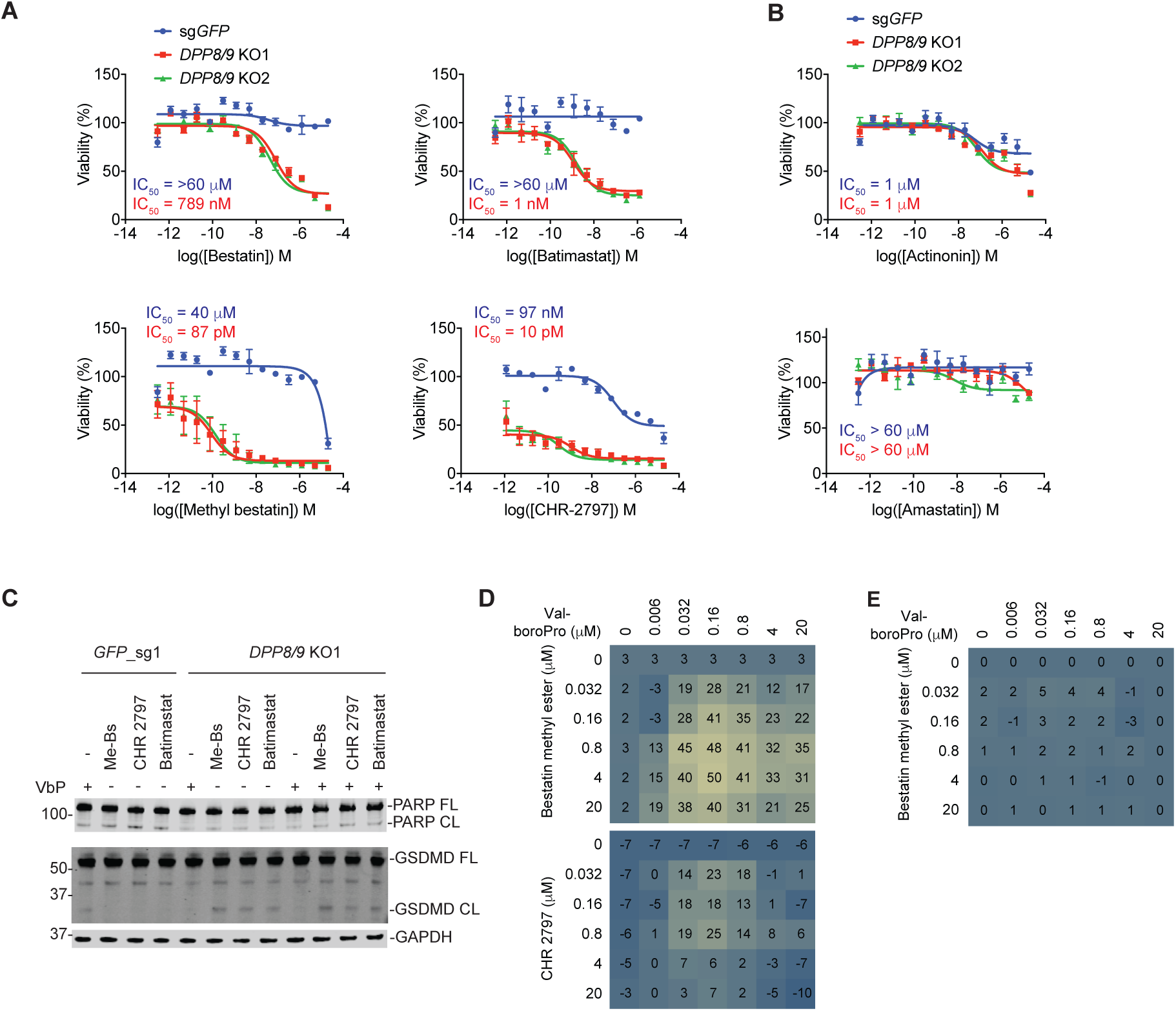
Aminopeptidase inhibition is synergistic with DPP8/9 inhibition in human THP-1 cells. (**A,B**) *DPP8/9* knockout and control THP-1 cells were treated with the indicated compounds for 48 h before cell viability was assessed by CellTiter-Glo. Bestatin, methyl bestatin, batimastat, and CHR-2797 preferentially killed the *DPP8/9* knockout cells (**A**), whereas actinonin and amastatin did not (**B**). Data are means ± SEM of three biological replicates. (**C**) *DPP8/9* knockout and control THP-1 cells were treated with the indicated compounds (2 μM, 24 h) before lysates were harvested and PARP cleavage (i.e., apoptosis) and GSDMD cleavage (i.e., pyroptosis) were evaluated by immunoblotting. No PARP cleavage was observed, but the aminopeptidase inhibitors induced GSDMD cleavage selectively in the *DPP8/9* knockout cells. VbP only induced GSDMD cleavage in control cells, as expected. (**D,E**) Control (**D**) or caspase-1 knockout (**E**) THP-1 cells were treated with the indicated compounds at the indicated concentrations for 24 h before cell viability was determined by CellTiter-Glo. For each pair of concentrations, we subtracted the predicted Bliss additive effect from the observed inhibition. Values greater than zero indicate synergy. We observed synergy in the THP-1 control cells with bestatin methyl ester and CHR 2797 (**D**), but not in the VbP-resistant *CASP1^-/-^* THP-1 cell line (**E**). Data are means ± SEM of three biological replicates.

**Table S1.**
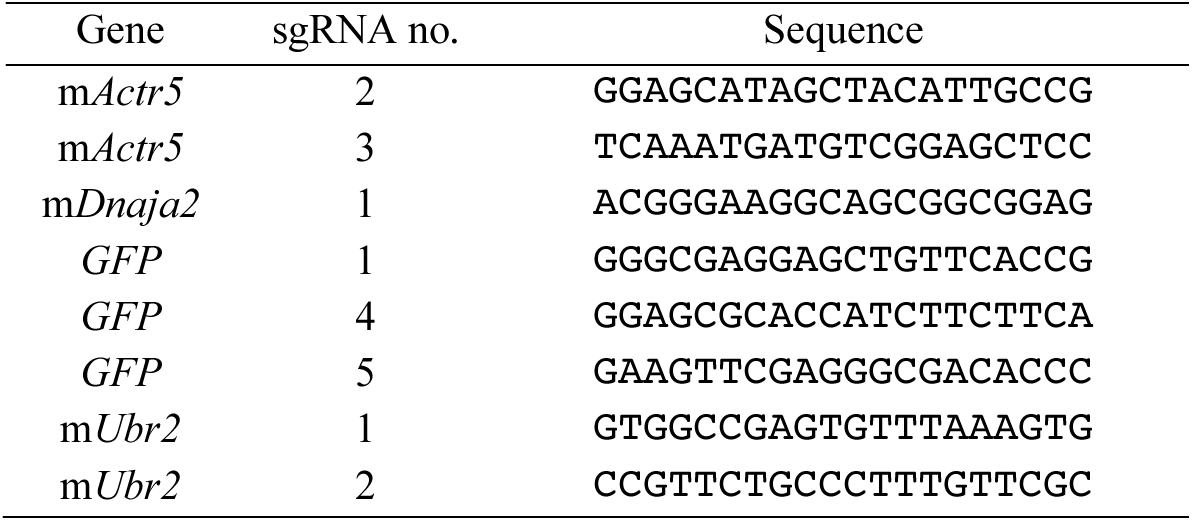
sgRNA sequences used in this study.

